# The cervical and meningeal lymphatic network as a pathway for retrograde nanoparticle transport to the brain

**DOI:** 10.1101/2024.01.06.574478

**Authors:** Héctor M Ramos-Zaldívar, Iva Polakovicova, Edison Salas-Huenuleo, Claudia P Yefi, David Silva, Pedro Jara-Guajardo, Juan Esteban Oyarzún, Álvaro Neira-Troncoso, Patricia V. Burgos, Viviana A. Cavieres, Eloisa Arias-Muñoz, Carlos Martínez, Ana L. Riveros, Alejandro H Corvalán, Marcelo J Kogan, Marcelo E Andia

**Author notes:** Corresponding author: Héctor M. Ramos-Zaldívar Contact information: Primary. Deceased author (Iva Polakovicova).

## Abstract

The meningeal lymphatic vessels have been described as a pathway that transports cerebrospinal fluid and interstitial fluid in a unidirectional manner towards the deep cervical lymph nodes. However, these vessels exhibit anatomical and molecular characteristics typical of initial lymphatic vessels, with the absence of surrounding smooth muscle and few or absent valves. Given its structure, this network could theoretically allow for bidirectional motion. Nevertheless, it has not been assessed as a potential route for nanoparticles to travel from peripheral tissues to the brain. Here we show that extracellular vesicles derived from the B16F10 melanoma cell line, along with superparamagnetic iron oxide nanoparticles, gold nanorods, and Chinese ink nanoparticles can reach the meningeal lymphatic vessels and the brain of C57BL/6 mice after administration within deep cervical lymph nodes *in vivo,* exclusively through lymphatic structures. Since the functional anatomy of dural lymphatics has been found to be conserved between mice and humans, we expect that our results will encourage further research into the retrograde motion of nanoparticles towards the brain for pharmacological purposes in nanomedicine, as well as to better understand the fluid dynamics in different physiological or neuropathological conditions.

## Introduction

The lymphatic system, including the meningeal lymphatic vessels, has usually been described as a unidirectional transport system of fluid and macromolecules from tissues to venous circulation^1,2^. This concept has prevented its examination as a nanoparticle and drug delivery pathway to the brain, as administered contents would be expected to all be cleared to the thorax. Nevertheless, evaluating nanoparticle flow through these vessels is relevant considering the participation of the lymphatic system in immune transportation, its pathologic involvement in cancer metastasis and the spreading of some infectious diseases, as well as its potential as a drug delivery pathway for its targeting and pharmacokinetic advantages, including bypassing first-pass metabolism in the liver^3^.

Studies in mice have described the anatomical and morphological characteristics of meningeal lymphatics to be consistent with initial lymphatic vessels^4,5^. This includes a noncontinuous basement membrane, sparse or no lymphatic valves, and no smooth muscle cell lining^4,5^; which implies that meningeal lymphatic vessels might not have a preferential flow determined by its own structural components. Together, these findings open a theoretical possibility for retrograde flow towards the brain that would depend on the physiological and mechanical conditions of the vessels.

Here, we suggest that the cervical and meningeal lymphatic system can transport nanoparticles not only towards the thorax but can also serve to carry particles towards the brain.

### MRI imaging of SPIONs and SPION-loaded exosomes in cervical and meningeal lymphatic vessels

Extracellular vesicles (EVs) are membranous particles naturally emitted by cells, encased in a lipid bilayer, and unable to undergo replication^6^. Exosomes are a subset of EVs that have an endosomal origin and a size range of 40 to 160 nm (average 100 nm)^7^. How EVs can travel between peripheral tissues and the brain in a bidirectional manner remains poorly understood^8^. To investigate if the cervical and meningeal lymphatic system is a possible route for EVs crossing to and from the central nervous system (CNS), we prepared exosomes loaded with superparamagnetic iron oxide nanoparticles (SPIONs) to evaluate their anterograde and retrograde directional flow through MRI imaging, leveraging the high efficiency of SPIONs as contrast agents^9^. The anterograde directional flow was defined as the classically described motion of lymphatic components towards the thorax following an injection into the cisterna magna. The retrograde directional flow was defined as nanoparticle motion towards the meningeal lymphatic vessels and the brain after a deep cervical lymph node injection.

We first prepared SPIONs through coprecipitation of ferric and ferrous chlorides with ammonium associated with an acidic pH. This produced SPIONs with mean diameter of 14.28 ± 5.57 nm measured by dynamic light scattering (DLS) (Fig. 1a). Scanning transmission electron microscopy (STEM) confirmed SPIONs mean size of 7.08 ± 2.2 nm (Fig. 1a). The zeta potential was positive at 36.9 ± 0.51 mV (Fig. 1a). After a ten-fold stock dilution with Milli-Q water, pH was increased to 7. Next, we proceeded to isolate exosomes from the B16F10 melanoma cell line using the Exo-spin (CELL GS) protocol. The MicroBCA (ThermoScientific) assay kit was used for total protein quantitation, yielding 300 μg/mL. Western blot analysis confirmed the presence of EV markers EEA1 and TSG101 (Fig. 1b). Finally, suspended exosomes were electroporated in 4 mm path length electroporation cuvettes. A single pulse was applied to each exosome sample under the high voltage setting and at an electric field of 0.75 kV/cm. After reisolating the labeled exosomes with Exo-spin columns, DLS revealed an exosome population of an average size of 106 nm ± 27.35 and a mean zeta potential of -17.1 ± 0.53 mV (Fig. 1c). Fig. 1c shows both B16F10 exosomes and exosomes electroporated with SPIONs as seen by electron microscopy.

**Fig. 1:**
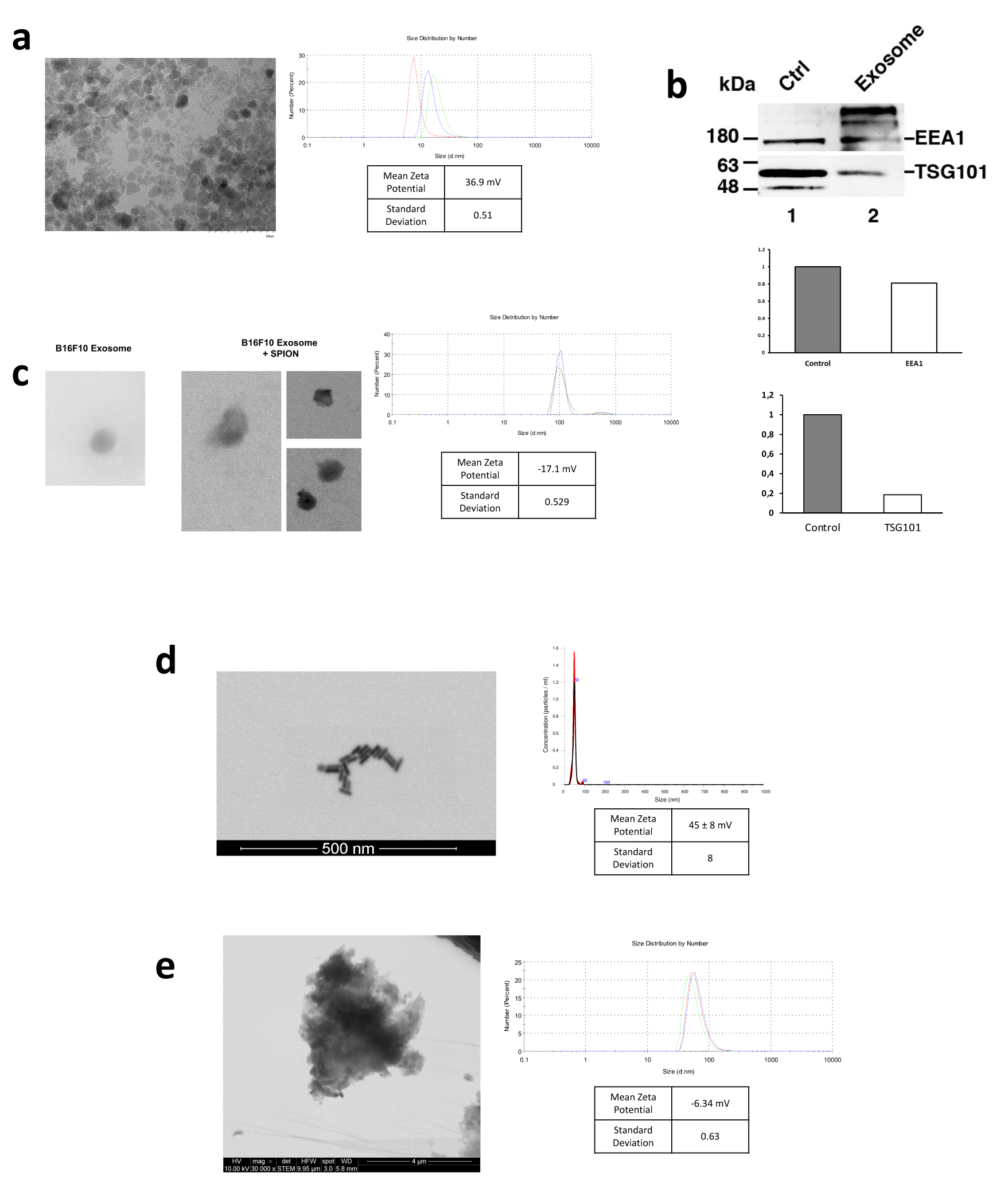
Characterization of nanoparticles. **a**, STEM visualization of SPIONs; size distribution and zeta potential measured by DLS. **b,** STEM visualization of exosomes with and without SPIONs labeling; size distribution and zeta potential measured by DLS. **c,** Western blot of EV markers EEA1 and TSG101 on exosomes from the B16F10 melanoma cell line. Control was performed using a cellular extract from the B16F10 cell line. Quantification of EEA1 and TSG101 with respect to control is shown. **d,** STEM visualization of gold nanorods; size distribution measured by NTA and zeta potential determined by DLS. **e,** STEM visualization of Chinese ink; size distribution and zeta potential measured by DLS.

To evaluate the retrograde directional flow through the cervical and meningeal lymphatic system, C57BL/6 mice (n=3 per condition) were injected *in vivo* with a 10 μL solution of either SPIONs (3200 μg/mL) or SPION-loaded exosomes (1.67 x 10^11^ particles/mL). These were compared with control mice with no injections (n=3). Animals were anesthetized with 5% isoflurane for 5 min and kept under anesthesia with a nasal cannula supplying 1%-2% isoflurane during the entire procedure. A syringe with a 30G needle was loaded with 10 μL of each solution and administered into the deep cervical lymph node. To locate the lymph node, skin and subcutaneous tissue were dissected at the midline of the neck, extending the field laterally at the supraclavicular area until both mandibular glands were exposed. Glands were detached from the clavicle surface and moved cranially. The sternocleidomastoid muscles were then displaced until the deep cervical nodes, surrounded by adipose tissue, were identified. Euthanasia by intraperitoneal sodium thiopental overdose (100 mg/kg) was performed 30 min after injection. The head and neck were preserved by fixation with 4% paraformaldehyde for MR imaging (Philips Achieva 1.5 T MR scanner) and histological analysis.

Both SPIONs and SPION-loaded exosomes revealed hypointense signals in the brain ventricles and parenchyma, particularly in the T2* MRI maps (Fig. 2a). Hypointense signals were also detected at the level of the neck where the injections were administered. Retrograde directional flow of SPIONs injected into the deep cervical lymph node *in vivo* was observed in staining of neck and head lymphatic vessels, including meningeal lymphatics, in all mice (n=3) (Supplementary Fig. S1 and Supplementary Fig. S2c). However, no staining was observed within the brain parenchyma with the Perls’ Prussian Blue technique. Iron detection through this staining method is prone to yield false negatives, as the detection requires the accumulation of several hundreds of nm in diameter^10^, which could hinder signals from SPIONs smaller than 10 nm diluted in the volume of the brain parenchyma. Combined Perls’ Prussian Blue staining and anti-LYVE-1 (lymphatic vessel endothelial hyaluronan receptor-1) immunohistochemistry revealed nanoparticles within the cervical lymphatic vessels towards the meningeal lymphatic vessels in the *in vivo* retrograde directional flow experiments performed on all C57BL/6 mice (n=3). No SPION staining was detected within arterial or venous structures within the head and neck (Supplementary Fig. S3b). As expected, exosomes loaded with SPIONs did not stain, indicating the presence of iron nanoparticles within exosome membranes.

**Fig. 2:**
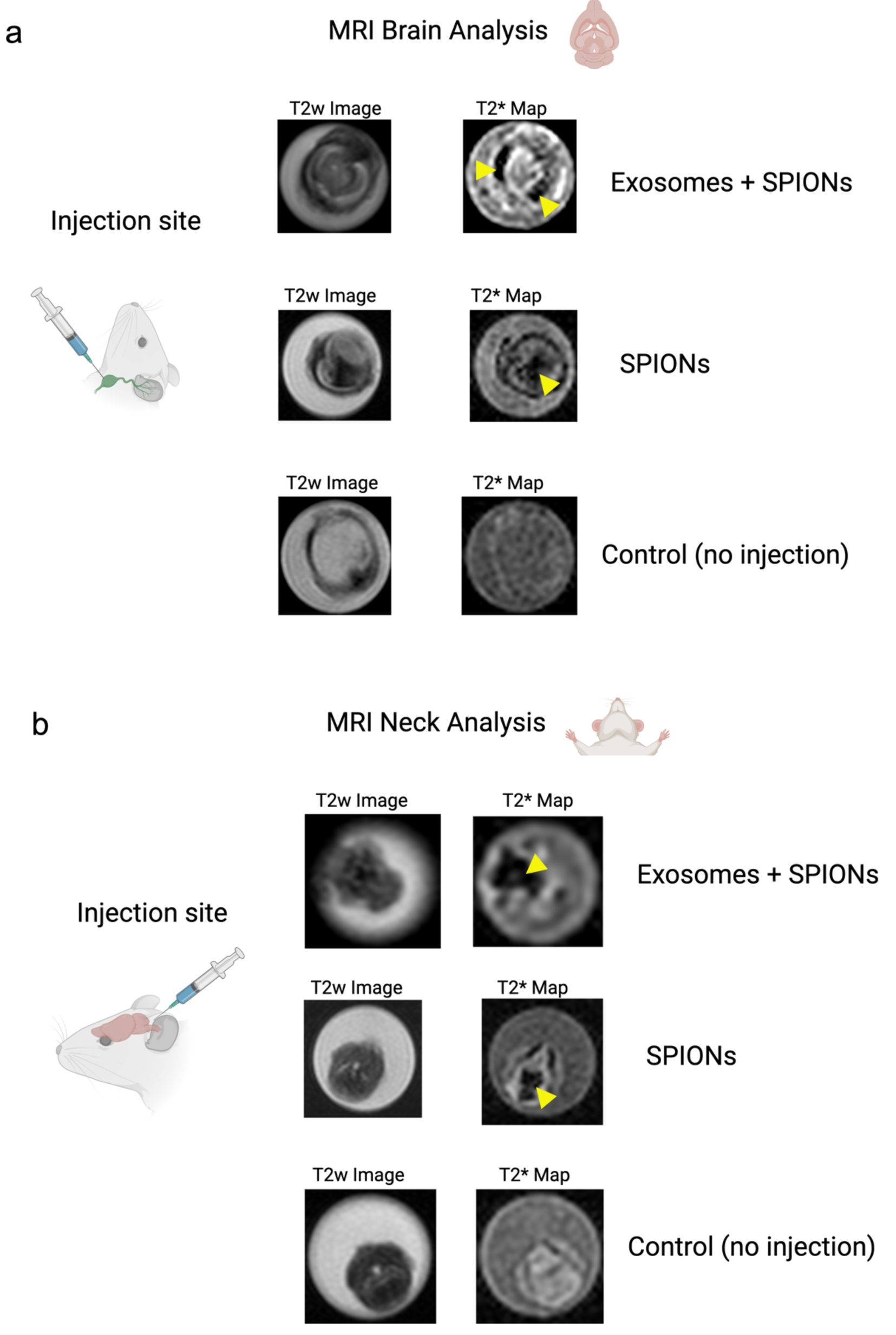
Directional flow analysis by MRI of SPIONs and SPION-labeled exosomes through the cervical and meningeal lymphatic system. **a**, Retrograde Directional Analysis: Brain images reveal the detection of nanoparticles in this region 30 minutes after injection into the deep cervical lymph node (n=3), particularly evident in the T2* map (yellow arrows). These two conditions were compared to control mice with no injected solutions (n=3). **b,** Anterograde Directional Analysis: Neck images reveal the detection of nanoparticles in this region 30 minutes after injection into the cisterna magna (n=3), particularly evident in the T2* map (yellow arrows). These two conditions were compared to control mice with no injected solutions (n=3).

To evaluate the anterograde directional flow, we injected C57BL/6 mice (n=3 per condition) *in vivo* with a 10 μL solution of either SPIONs (3200 μg/mL) or SPION-loaded exosomes (1.67 x 1011 particles/mL). These were compared with control mice with no injections. Animals were anesthetized with 5% isoflurane and kept under anesthesia with a nasal cannula supplying 1% isoflurane during the entire procedure. A syringe with a 30G needle was loaded with 10 μL of each solution and administered into the cisterna magna; by placing the mouse in prone position, flexing the head at a 135° angle with the body, and penetrating directly underneath and laterally to the end of the occipital bone towards the foramen magnum through the intact skin. Euthanasia by intraperitoneal sodium thiopental overdose (100 mg/kg) was performed 30 min after injection. The head and neck were preserved by fixation with 4% paraformaldehyde for 1.5 Tesla MRI scanner and histological analysis.

Both SPIONs and SPION-loaded exosomes showed hypointense signals of cervical lymphatic structures after intracerebroventricular injections through the cisterna magna, as seen in the T2w images and T2* maps (Fig. 2b). Anterograde directional flow in *in vivo* procedures (n=3) of SPIONs after administration into the cisterna magna was confirmed by the detection of Perls’ Prussian Blue staining in cervical lymphatic vessels in all mice (Supplementary Fig. S4). At the level of the head, the injection into the cisterna magna also showed staining within the ventricles. As expected, exosomes loaded with SPIONs did not stain, indicating the presence of iron nanoparticles within exosome membranes.

Together, MRI imaging results indicate that the cervical and meningeal lymphatic system can transport SPIONs and SPION-labeled exosomes both towards the thorax and in the direction of the brain.

### Gold nanorods bidirectional motion through the cervical and meningeal lymphatic system

With MRI imaging suggesting the possibility of retrograde flow of SPIONs and SPION-loaded exosomes towards the brain after a cervical administration, we further examined other nanoparticles that could be more effectively assessed through histological techniques in the brain parenchyma. This also allowed the evaluation of different alternatives that can be subsequently explored in pharmacology and nanomedicine. Therefore, we used gold nanorods functionalized with polyethylene glycol (GNR-PEG) as described in Methods. GNR-PEG with a mean size of 49.1 ± 0.9 nm and mean zeta potential of 45 ± 8 mV were obtained and measured by DLS and Nanoparticle Tracking Analysis (NTA) (Fig. 1d). GNR-PEG morphology can be observed in the STEM image seen in Fig. 1d. The GNR-PEG size distribution determined by STEM showed a length of 34.6 ± 4.2 nm and a width of 11.4 ± 1.6 nm. The anterograde and retrograde directional flow were evaluated in two scenarios with *post-mortem* or *in vivo* administrations of GNR-PEG solutions in C57BL/6 mice (n=3 per condition) into the cisterna magna or the deep cervical lymph node. These interventions were compared with control mice with no injections (n=3).

*Post-mortem* procedures were performed using 50 μL of GNR-PEG at a concentration of 1.71 x 10^14^ particles/mL. Dissections for identifying the deep cervical lymph node during retrograde administrations followed the method described previously for *in vivo* procedures. The head and neck were preserved by fixation with 4% paraformaldehyde for histological Gold Enhancement (Nanoprobes GoldEnhance TM LM Kit) analysis. Retrograde directional flow in *post-mortem* procedures (n=3) of gold nanorods after deep cervical lymph node administration was confirmed by the detection of Gold Enhancement staining at different CNS regions in all mice. These included the olfactory bulb, the brain parenchyma, and within the meningeal lymphatic vessels (Fig. 3a). No staining was detected within arterial or venous structures within the head and neck, ruling out other sources of nanoparticle distribution to the brain in *post- mortem* GNR-PEG assays. Control mice with no GNR-PEG administrations showed no Gold Enhancement staining in any anatomical structure.

**Fig. 3:**
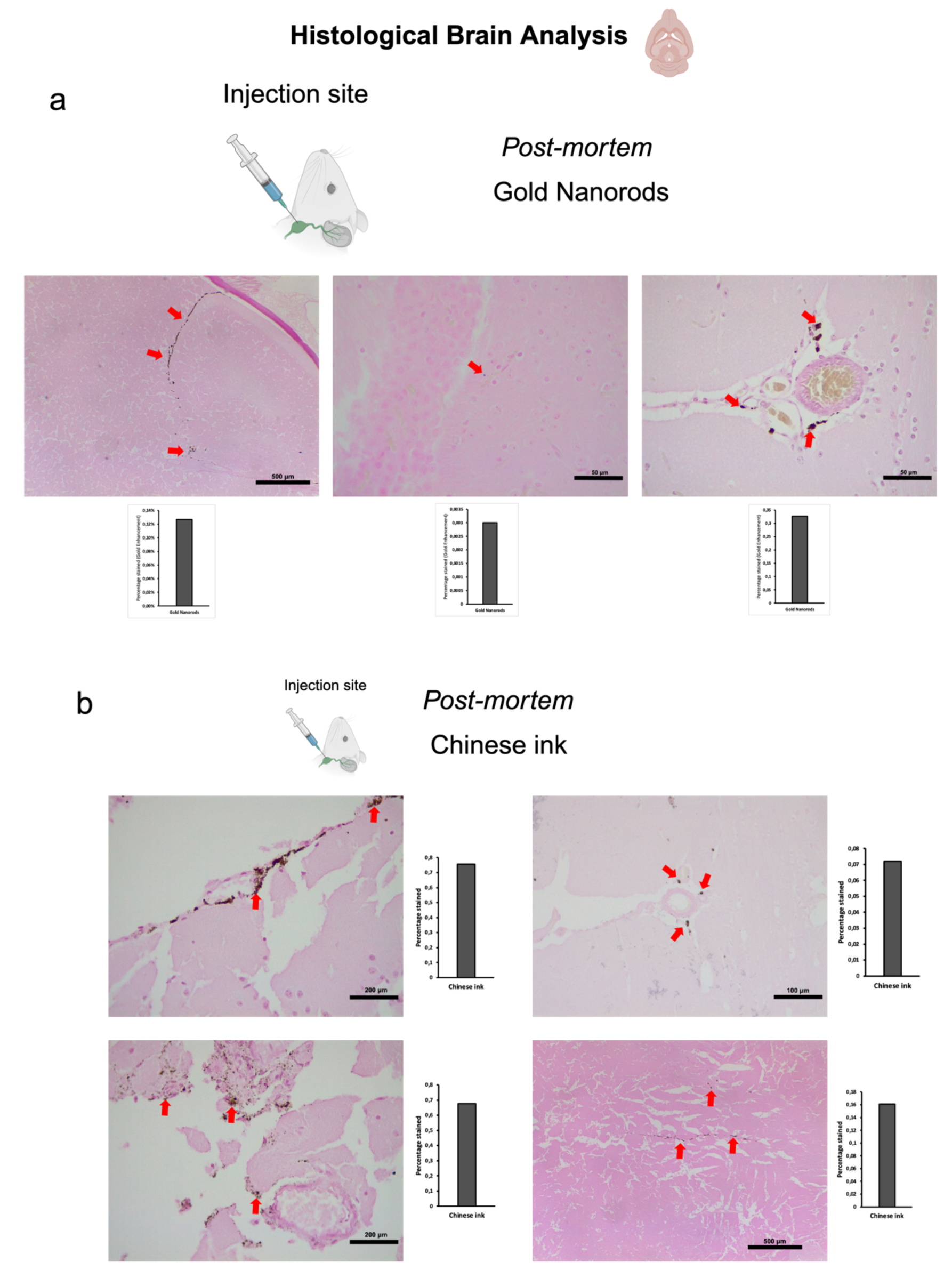
Retrograde directional flow analysis by brain histology after *post-mortem* nanoparticle administration into the deep cervical lymph node. **a**, Gold nanorods were identified by the Gold Enhancement technique in the olfactory bulb, the brain parenchyma, and the meningeal lymphatic vessels (red arrows) (n=3). **b,** Chinese ink nanoparticles stained the meningeal lymphatic vessels, the brain parenchyma, and the third ventricle wall (red arrows) (n=3).

Anterograde directional flow in *post-mortem* procedures (n=3) of GNR-PEG after administration into the cisterna magna was also confirmed by the detection of Gold Enhancement staining in cervical lymphatic vessels in all mice, as well as in the connective tissue of the neck (Fig. 5a). Gold nanoparticles were also identified in the cervical spinal cord as well as its surrounding subdural space and associated peripheral nerves (Fig. 5a). At the level of the head, the injection into the cisterna magna also showed staining within lateral ventricles, the third ventricle, the olfactory bulb, and the optic chiasm in all mice. No Gold Enhancement staining was observed in any anatomical structure of the control mice that did not receive GNR-PEG administrations.

To investigate the directional flow of GNR-PEG under physiological conditions and to minimize the potential impact of volume, we administered 10 μL of GNR-PEG at a concentration of 1.71 x 10^14^ particles/mL *in vivo*. The administration technique was as previously described for deep cervical lymph node and cisterna magna injections. Combined Gold Enhancement and anti-LYVE-1 immunohistochemistry revealed gold nanoparticles within the cervical and meningeal lymphatic vessels in the *in vivo* retrograde directional flow experiments performed on all C57BL/6 mice (n=3) (Fig. 4a). GNR-PEG also reached the brain parenchyma through the retrograde flow from the cervical lymphatic vessels in all mice (Fig. 4a). GNR-PEG were found staining within anti-LYVE1 cervical lymphatic vessels towards the meningeal lymphatics (Supplementary Fig. S2a). No staining was detected within arterial or venous structures within the head and neck, ruling out other sources of nanoparticle distribution to the brain in *in vivo* gold nanoparticle assays (Supplementary Fig. S3a). Anterograde directional flow of gold nanoparticles after administration into the cisterna magna *in vivo* was established by the detection of Gold Enhancement staining in cervical lymphatic vessels in all mice (Fig. 6a). Gold Enhancement staining was not detected in any anatomical structure of the control mice that did not undergo GNR-PEG administrations.

**Fig. 4:**
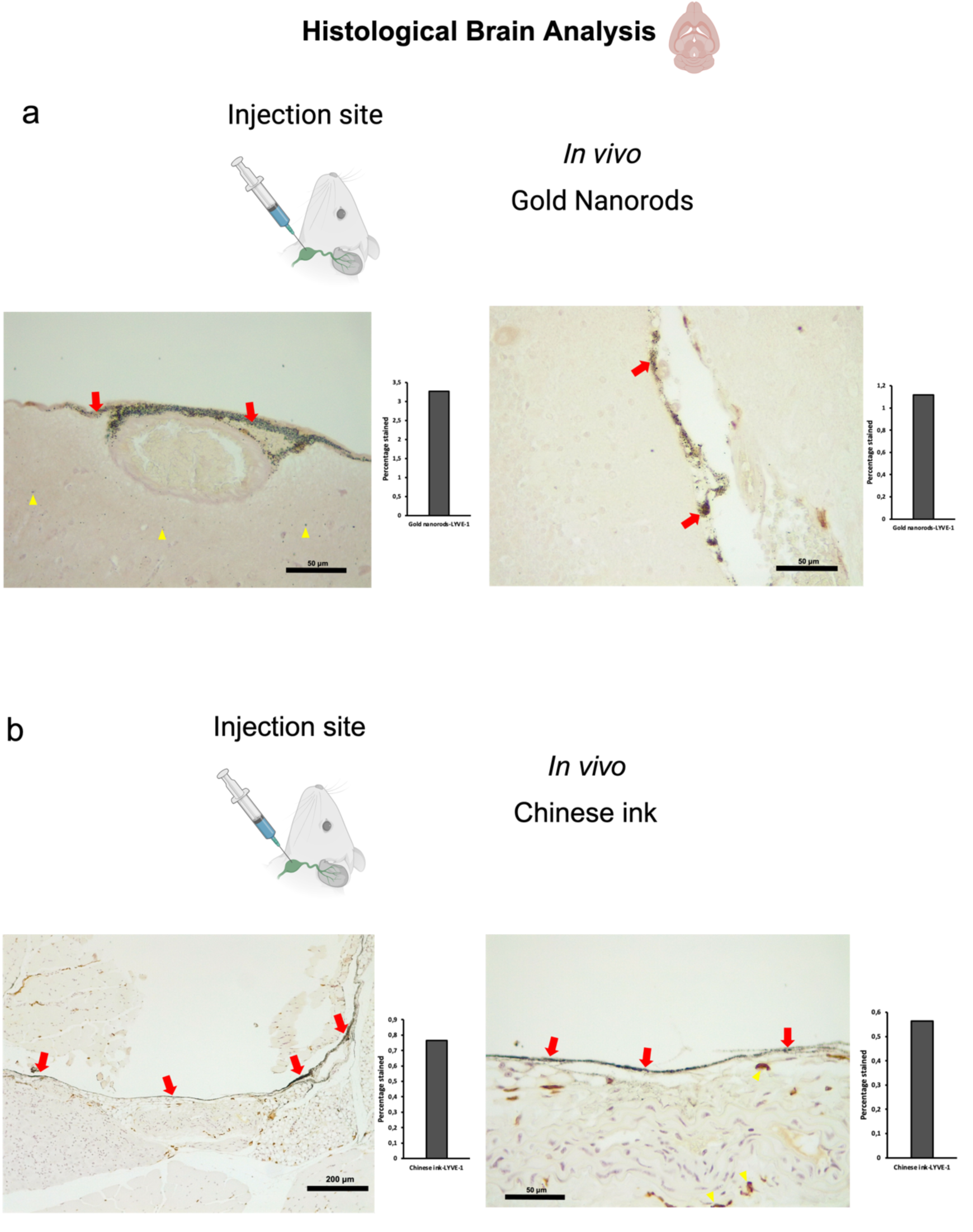
Retrograde directional flow analysis by brain histology after *in vivo* nanoparticle administration into the deep cervical lymph node. **a**, Combined Gold Enhancement and anti-LYVE-1 immunohistochemistry showed gold nanorods within meningeal lymphatic vessels (red arrows) and the brain parenchyma (yellow arrows), with no staining within cerebral arteries (n=3). **b,** Meningeal lymphatic vessels stained with anti-LYVE-1 immunohistochemistry and colocalized with Chinese ink nanoparticles (red arrows). Chinese ink was also identified in the brain parenchyma (yellow arrows) (n=4). LYVE-1: lymphatic vessel endothelial hyaluronan receptor-1.

**Fig. 5:**
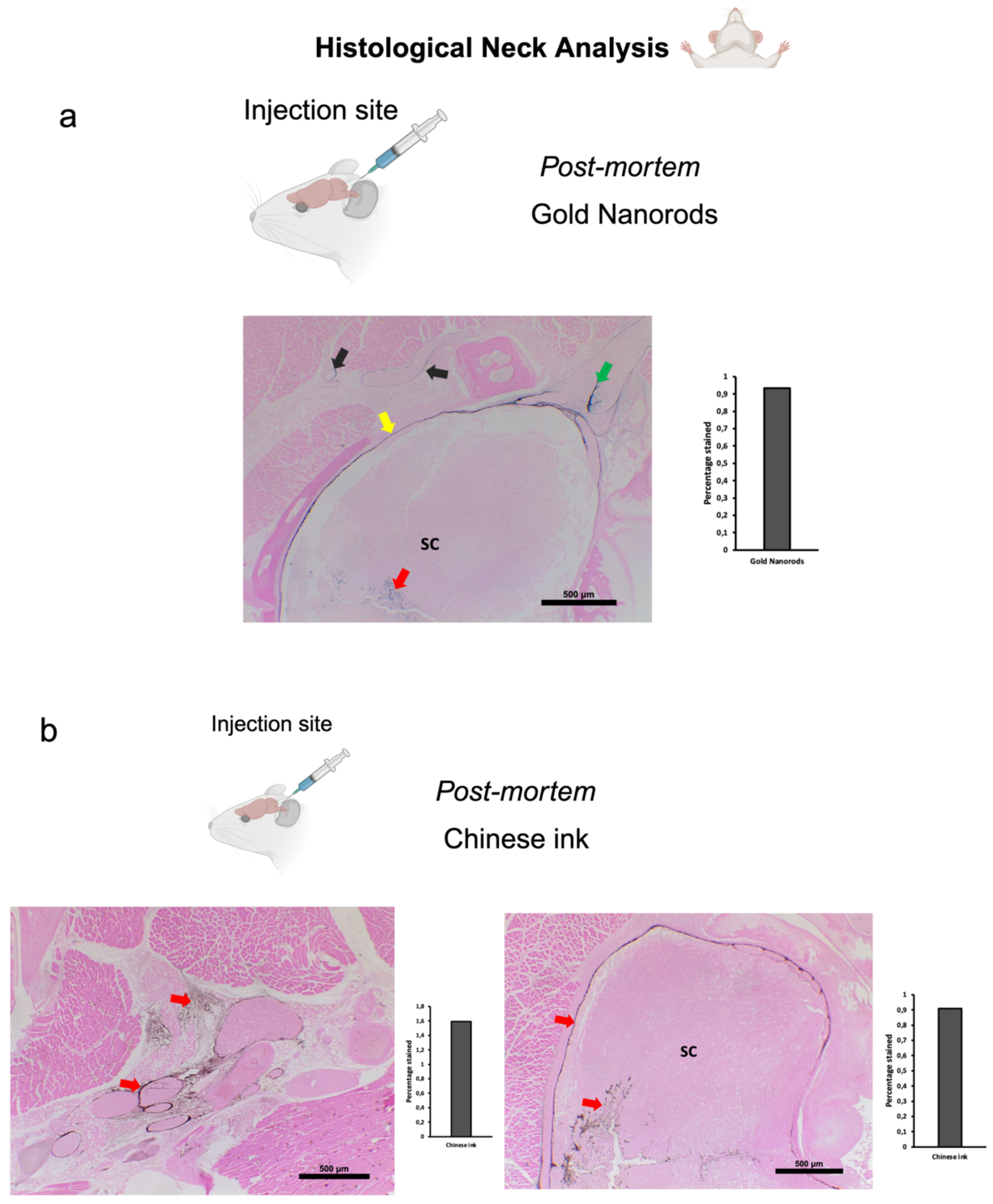
Anterograde directional flow analysis by brain histology after *post-mortem* nanoparticle administration into the cisterna magna. **a**, Gold Enhancement showed staining of the cervical spinal cord (red arrow), its surrounding subdural space (yellow arrow) and associated peripheral nerves (green arrow). Gold nanoparticles were also detected in cervical lymphatic vessels and connective tissue (black arrows) (n=3). **b,** Chinese ink nanoparticles were identified in cervical lymphatic vessels, connective tissue, as well as the cervical spinal cord and its surrounding subdural space (indicated by red arrows) (n=3). SC: spinal cord.

The results of histological analyses collectively suggest that the cervical and meningeal lymphatic system is capable of bidirectional transportation of gold nanorods, encompassing movement towards both the thorax and the brain.

### Chinese ink bidirectional motion through the cervical and meningeal lymphatic system

Chinese ink has been previously used for staining of lymphatic structures^11–13^, which led to its consideration for our evaluation on the bidirectional motion of nanoparticles through the cervical and meningeal lymphatic system. Interestingly, Chinese ink (Artel, Santiago, Chile) was characterized by DLS, which revealed nanoparticles with a mean (± SD) size of 61.62 ± 4.84 nm and mean (± SD) surface zeta potential of -6.34 ± 0.63 mV (Fig. 1e), which are measurements similar to the range values observed in purified exosomes. To our surprise, we found that Chinese ink can also be stained with the Gold Enhancement technique (Nanoprobes GoldEnhance TM LM Kit) used previously with GNR, confirmed by the lack of staining of control mice brain parenchyma slides with no nanoparticle administration. The anterograde and retrograde directional flow were evaluated after *post-mortem* and *in vivo* administrations of Chinese ink solutions in C57BL/6 mice (n=3 per condition) in the cisterna magna and the deep cervical lymph node, respectively. These were compared with control mice with no injections (n=3).

*Post-mortem* procedures were performed using 50 μL of 10% Chinese ink (Artel, Santiago, Chile). Dissections for identifying the deep cervical lymph node during retrograde administrations followed the method described previously for procedures with SPION-loaded exosomes. The head and neck were preserved by fixation with 4% paraformaldehyde for histological Gold Enhancement (Nanoprobes GoldEnhance TM LM Kit) analysis. Retrograde directional flow in *post-mortem* procedures (n=3) of Chinese ink after deep cervical lymph node administration was confirmed by the detection of Gold Enhancement staining at different CNS regions in all mice (Fig. 3b). Staining was detected within the meningeal lymphatic vessels, the third ventricle, and cortical regions near meningeal lymphatic vessels. No staining was detected within arterial or venous structures within the head and neck, ruling out other sources of nanoparticle distribution to the brain in *post-mortem* Chinese ink assays. In control mice that did not undergo Chinese ink administrations, there was no presence of Gold Enhancement staining in any anatomical structure.

Anterograde directional flow in post-mortem procedures (n=3) of Chinese ink nanoparticles after administration into the cisterna magna was confirmed by the detection of staining in cervical lymphatic vessels in all mice, as well as in connective tissue of the neck (Fig. 5b). Chinese ink nanoparticles were also identified in the cervical spinal cord as well as its surrounding subdural space (Fig. 5b). At the level of the head, the injection into the cisterna magna also showed staining within lateral ventricles. The anatomical structures of the control mice that did not receive Chinese ink administrations exhibited no signs of Gold Enhancement staining.

To examine the Chinese ink directional flow under physiological conditions and reducing the potential effects of volume, we conducted *in vivo* administrations of 10 μL of 10% Chinese ink (Artel, Santiago de Chile). The administration procedure followed the previously established method for injections into the deep cervical lymph nodes and the cisterna magna. Retrograde directional flow of *in vivo* procedures (n=4) was confirmed in all mice after deep cervical lymph node administration. Combined Gold Enhancement and anti-LYVE-1 immunohistochemistry showed Chinese ink within the meningeal lymphatic vessels and cortical regions near these lymphatic structures (Fig. 4b). One mouse died at minute two before the expected completion time of 30 min before euthanasia. Nevertheless, after histological analysis of this specimen, Chinese ink nanoparticles were identified in the meningeal lymphatic vessels and the brain parenchyma. Chinese ink was found staining within anti-LYVE1 cervical lymphatic vessels towards the meningeal lymphatics (Supplementary Fig. S2b). No staining was detected within arterial or venous structures within the head and neck of two out of three mice (Supplementary Fig. S3c). One mouse presented staining within the jugular vein but not the carotid artery (Supplementary Fig. S3), which also indicates that the observed nanoparticles at the meningeal lymphatic vessels and the brain parenchyma originate mainly from the lymphatic system distribution and not through arterial circulation of cardiac and other thoracic vessels. Anterograde directional flow in *in vivo* procedures (n=3) after administration into the cisterna magna was confirmed by the detection of staining in cervical lymphatic vessels in all mice (Fig. 6b). Chinese ink nanoparticles were also identified in the subarachnoid space, the cervical spinal cord, and peripheral nerves (Fig. 6b). No staining was observed in any anatomical structure of the control mice that did not receive any administrations.

**Fig. 6:**
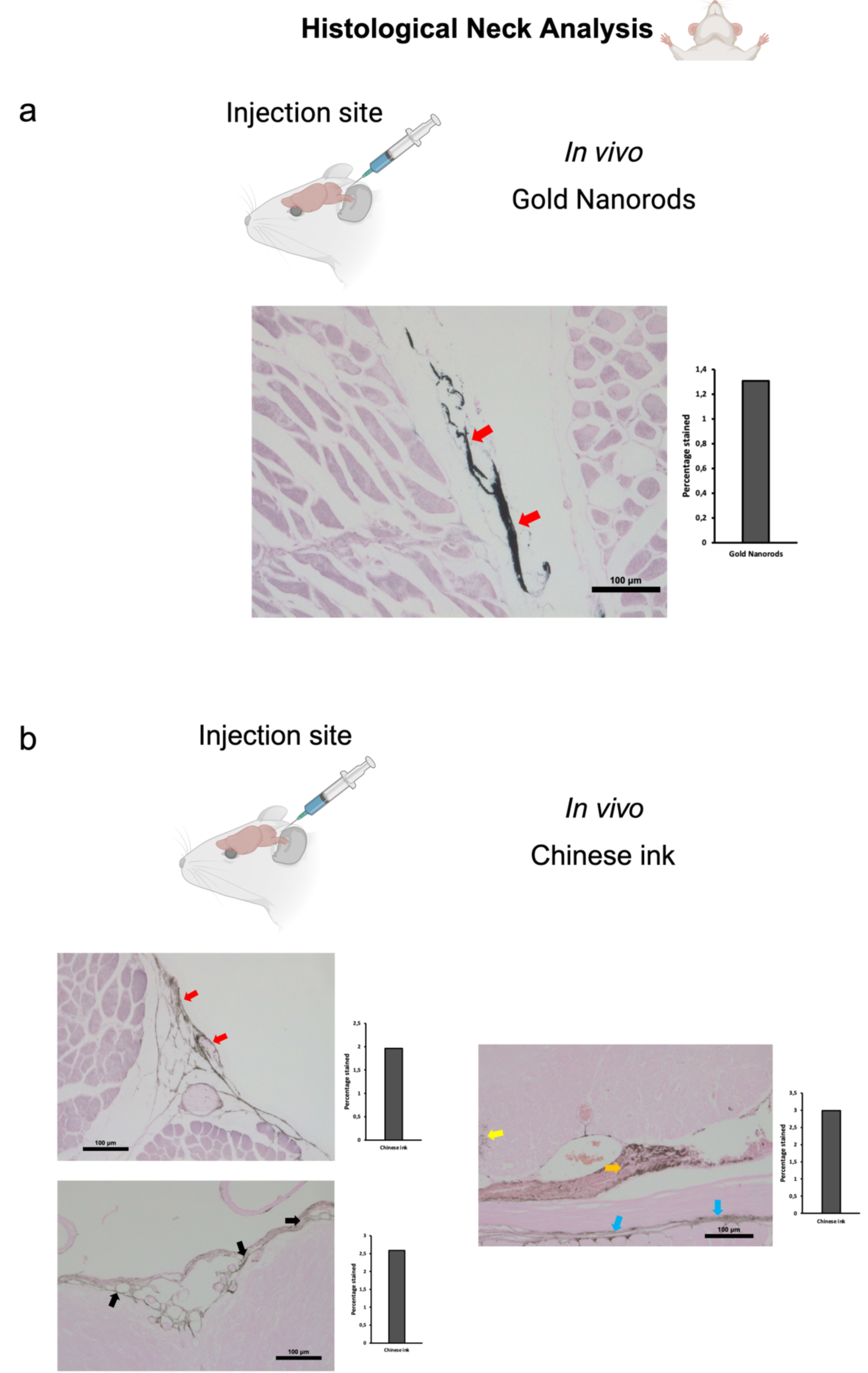
Anterograde directional flow analysis by brain histology after *in vivo* nanoparticle administration into the cisterna magna. **a**, Gold Enhancement showed staining of lymphatic vessels in the cervical region (red arrows) (n=3). **b,** Chinese ink nanoparticles were identified in the cervical lymphatic vessels (red arrows), the subarachnoid space (black arrows), as well as the cervical spinal cord (yellow arrow), peripheral nerves (orange arrow), and connective tissue (blue arrows) (n=3).

Taken together, the findings of histological analyses indicate that the cervical and meningeal lymphatic system is capable of bidirectional transportation of Chinese ink nanoparticles, involving movement towards both the thorax and the brain.

## Discussion

We have shown evidence that suggests that the cervical and meningeal lymphatic system can transport nanoparticles not only in the classically described lymphatic drainage towards the thorax but can also serve as an access gate to the brain. SPIONs and SPION-loaded exosomes were detected by MRI in the brain of C57BL/6 mice after deep cervical lymph node administration *in vivo*. Gold nanorods and Chinese ink nanoparticles were also identified within the meningeal lymphatic vessels and the brain parenchyma of mice in the retrograde directional flow histological analysis from cervical injections in *post-mortem* and *in vivo* procedures. Anterograde directional flow experiments from all nanoparticle experiments also showed motion from the cisterna magna to the deep cervical lymph nodes. Together, these indicate that the system allows for bidirectional flow after administration.

Two pertinent factors to examine regarding this newly described retrograde lymphatic flow towards the brain include alternative vascular pathways and the influence of pressure exerted at the cervical injection site. The initial consideration, particularly in *in vivo* assays, was whether nanoparticles in meningeal lymphatic vessels and the brain might have originated from their distribution from the cervical lymphatic vessels, passing through the jugular vein, the superior vena cava, the right atrium and ventricle of the heart, through the pulmonary circulation, to the left atrium and ventricle of the heart, and towards the carotid arteries before entering the cerebral circulation. This would entail that nanoparticles would have been present in histological analyses within the jugular veins and the common carotid arteries during cervical examinations. However, as previously indicated, all *in vivo* SPION, GNR-PEG, and Chinese ink retrograde experiments (collectively n=10) showed no staining of carotid arteries. Moreover, nine out of the ten deep cervical lymphatic nanoparticle administrations showed no staining of jugular veins. GNR-PEG and Chinese ink post-mortem retrograde injections (collectively n=6) also concurred with these findings. Given that nanoparticles were indeed found staining within anti-LYVE1 cervical lymphatic vessels towards the meningeal lymphatics, this data supports the conclusion that nanoparticles reaching the brain were following a lymphatic pathway.

It is important to consider that although the volume injected in *in vivo* procedures was small (10 μL) the pressure exerted at the cervical injection site could have been significantly greater than the intranodal pressure, changing the fluid dynamics within the lymph node. However, this relationship is complex to determine at this stage because cervical intranodal pressures have not yet been established. Previous studies have analyzed intranodal pressure of other anatomical locations in mice with different results. Bouta *et al.* described intranodal popliteal and axillary pressures of an average 9 and 12 cmH2O, respectively, in normal wild type mice^14^. Kato *et al.* and Miura *et al.* used MXH10/Mo-lpr/lpr, a mouse model that develops systemic swelling of lymph nodes, obtaining lower values for intranodal pressures^15,16^. Miura *et al.* showed that mean pressures within subiliac and axillary nodes were 0.10 cm H2O and 0.03 cm H2O, respectively^16^. When examining their data, Kato *et al.* found subiliac lymph nodes with a mean pressure of 1.63 cm H2O^15^. Finally, Rhoner *et al.* have even described subatmospheric pressures of -1 cm H2O in axillary and brachial lymph nodes of immune-competent C57Bl/6 mice^17^.

Another factor to consider is that the pressure that can be produced for any syringe at a predetermined speed depends on the force applied divided by the surface area of the syringe plunger^18^. While humans can apply considerable forces to a plunger, with an average maximum force of 79N, this implies that a lower injection force is needed to generate equivalent pressures when syringe caliber is reduced^18^. Therefore, injections performed by a human operator may differ considerably in force and pressure exerted depending on the syringe caliber and intranodal state. Future research should address cervical intranodal pressures and the biomechanics of fluid administration to better understand the fluid dynamics of this anatomical region.

It should be highlighted that mice and human functional anatomy of dural lymphatics has been found to be conserved. Jacob *et al.* found similar circa-cerebral meningeal lymphatic architecture and relationship with dural venous sinuses, with limited connections with the nasal lymphatic bed, and a conserved pattern of cavernous sinus associated vessels penetrating the skull through several bilateral foramina of the skull base^19^. They emphasized that murine models are relevant to predict the pathophysiological contribution of the dural lymphatic system and test lymphatic-targeted drugs in neurological disease models^19^. Here we evaluated different nanoparticles with pharmacological applications through these lymphatic vessels and point to a bidirectional potential that opens the possibility of a new access to the brain. Our experiments therefore can also give insights into possible human fluid dynamics that should be explored.

The exact drivers of this bidirectional flow could involve pressure changes within the lymphatic vessels and nodes in a system with few valves and no smooth muscle cell lining when entering the head. This means that anatomical position changes or physiological changes in pressure surrounding lymphatic tissues could create conditions favoring motion in the retrograde or anterograde flow when necessary. Pathological conditions producing pressure changes in or around lymphatic tissues could also promote and determine directional flow in the cervical and meningeal lymphatic system. Particularly in mammals such as humans that experience radical changes in head and neck dynamics with the upright and recumbent position, the possibility of bidirectional lymphatic flow could be relevant in many physiological processes such as during sleep.

The pharmacological implications of our findings could be important in the field of nanomedicine. The methodology used for labeling exosomes in this project could be modified to carry drugs through the lymphatic system and improve specific distribution to the brain. Further studies can evaluate if an interstitial injection in the neck could deliver enough nanoparticles through this system to develop noninvasive treatment procedures, as homing characteristics to lymph nodes have been seen in previous studies of SPION-loaded exosomes to popliteal lymphatics^20^. Given that cancer exosomes could potentially move towards the brain through this pathway, and that an anecdotal case report has suggested that cancer cells can move in a retrograde lymphatic manner in other organs even in valve-equipped lymphatic vessels^21^, mortality in these patients would be substantially reduced if cerebral metastatic mechanisms could be prevented. In this regard, future investigations can delve into cancer exosome lymphatic inhibitors, by regulating or blocking movement through these vessels.

In neurodegenerative diseases, the use of promising peptide inhibitors of polyglutamine aggregation (QBP1, NT17, and PGQ9P2) in Huntington’s disease has been hindered precisely because of poor BBB penetration and low bioavailability^22^. A cervical lymphatic route could be an attractive pathway to evaluate more efficient means for accessing the brain without complex nanoparticle constructions. A recent publication by Dominy et al. has associated *P. gingivalis* with Alzheimer’s disease^23^. Bacterial DNA and RNA found in the brain of patients with this disease could be transported by exosomes through the lymphatic system. Other rapidly rising fields, such as the connection between the gut microbiota with diseases such as autism, neurological disorders like multiple sclerosis, and mental disorders^24^, could potentially involve retrograde lymphatic flow of exosomes and different nanoparticles towards the brain.

In conclusion, the cervical and meningeal lymphatic system can serve as an access route for nanoparticles to the brain, allowing bidirectional flow. This newly discovered mechanism for the meningeal lymphatic pathway could be exploited in the theranostic field of nanomedicine to deliver drugs for the treatment of various neurological diseases. Additionally, our findings using exosomes from the metastatic B16F10 melanoma cell line could aid in a more profound comprehension of brain metastasis pathophysiology regarding the participation of extracellular vesicles.

## Methods

### Superparamagnetic iron oxide nanoparticle synthesis

Samples of iron oxide nanoparticles were prepared by a chemical coprecipitation process from FeCl3 ·6H2O (432 mg), and FeCl2 ·4H2O (159mg). Ferric and ferrous chlorides were dissolved in 19mL of Milli-Q water with vigorous magnetic agitation at room temperature. One mL of ammonium (25%) was added to the solution with vigorous magnetic agitation for 10 min. Then, three washes with Milli-Q water were performed, maintaining iron-nanoparticles in an 80 mL beaker with a neodymium magnet. Subsequently, the superparamagnetic iron oxide nanoparticles were washed twice with nitric acid. Finally, iron-nanoparticles were dissolved in Milli-Q water for later characterization. A ten-fold stock dilution with Milli-Q water increased pH to 7.

### Characterization of iron oxide nanoparticles

The morphology and particle size of SPIONs was investigated by scanning transmission electron microscopy (STEM, FEI Quanta 250) operating at 10.00 kV. The Malvern Zetasizer was used for dynamic light scattering size determination and superficial charge. SPIONs concentrations obtained from the synthesis were measured using Nanoparticle Tracking Analysis by NanoSight. pH of the solutions was determined by pH meter. The concentration of iron nanoparticles in solutions was determined by inductively coupled plasma mass spectrometry (ICP MS).

### Cell culture and exosome purification

The B16F10 melanoma cell line was cultured using Exo-free medium. The isolated supernatant was centrifuged twice: first at 300g for 10 min at 4 °C and then at 16000g for 20 min at 4 °C. Filtration followed through 0.2 μm pore size filters. For purification, an Exo-spin (CELL GS) protocol was conducted. Extracellular vesicles were first concentrated using a 10 kDa filter to separate larger cellular structures and then diluted in filtered PBS (0.1 μm pore size). The samples were then precipitated using Exo-spin Buffer overnight and then centrifuged for 1 hour at 16000g. The obtained pellet with exosomes was resuspended in 100 μL of PBS. Exo-spin columns were prepared with two consecutive washes with 250 μL of PBS at 50g for 10 sec. Finally, diluted exosomes were passed through the column using 200 μL of PBS and collected in microcentrifuge tubes.

### Nanoparticle internalization to exosomes

Suspended exosomes, purified as previously described, were electroporated in 4 mm path length electroporation cuvettes. A single pulse was applied to each exosome sample under the high voltage setting and at an electric field of 0.75 kV/cm. Following electroporation, nanoparticle-loaded exosomes were reisolated using the Exo-spin protocol.

### Characterization of exosomes

Exosomes were characterized by protein concentration (microBCA assay), shape by scanning transmission electron microscopy (STEM, FEI Quanta 250), concentration and size with NanoSight. Western blot analysis was performed to determine the presence of exosome markers EEA1 and TSG101. Cell extracts and exosomes were lysed at 4°C in lysis buffer (50 mM Tris-HCl pH 7.4, 150 mM NaCl, 1 mM EDTA, 1% Triton X-100) supplemented with a cocktail of protease inhibitors [416 μM 4-(2- Aminoethyl)benzenesulfonyl fluoride, 0.32 μM Aprotinin, 16 μM Bestatin, 5.6 μM E-64, 8 μM Leupeptin and 6 μM Pepstatin A; Sigma-Aldrich] and phosphatase inhibitors (1 mM NaF, 0,3 mM Na2P2O7 and 1 mM Na3VO4; Sigma-Aldrich). Cell lysates were collected and lysed for 30 min at 4°C in rotation. Extracts were further centrifuged for 20 min at 13.000xg at 4°C. Samples with an equivalent amount of protein were denatured at 65°C for 5 min with Laemmli SDS- PAGE sample buffer and analyzed by SDS-PAGE.

### Gold nanorods synthesis

For the preparation of a seed solution of gold nanoparticles, a cold-prepared sodium borohydride solution (600 μL, 0.01 M) was added to 250 μL of 0.01 M HAuCl4 in 9.75 mL of 0.1 M cetyltrimethylammonium bromide (CTAB) in a flask, under vigorous magnetic stirring. The seed solution was kept at 27 °C for 2 h, before use. After that, 55 μL of 0.1 M ascorbic acid solution (Sigma Chemical Co., St. Louis, MO, USA) was added to a growth solution containing 75 μL of 0.01 M AgNO3 (Sigma Chemical Co., St. Louis, MO, USA), 9.5 mL of 0.1 M CTAB, and 500 μL of 0.01 M HAuCl4. Further, 250 μL of 0.1 M HCl and 12 μL of the previously prepared seed solution were added. The solution was incubated for 10 min at 27 °C and then centrifuged at a 7030g for 15 min. After centrifugation, the supernatant was removed, and the pellet was resuspended in milli-Q water.

The GNRs were conjugated with asymmetrical PEGs that have a thiol group (SH) at one end, and a methoxy (HS-PEG-OMe MW 5K, JenKem Technology, TX, USA) or a carboxylic acid group (HS-PEG-COOH MW 5K, JenKem Technology, TX, USA) at the other. A total of 50 μL of 1 mM HS-PEG-OMe in a water solution was added to 10 mL of 1 nM GNRs-CTAB and stirred for 10 min. After centrifugation at RCF of 16,100g for 10 min, the pellet was resuspended in 10 mL of milli-Q water. Subsequently, 300 μL of 1 mM HS-PEG-COOH solution was added into the water solution, and the suspension obtained was stirred for one hour. Further, the suspension was centrifuged at 16,100g for 10 min, and the pellet was resuspended in 100 μL of 0.1 M 2-(N-morpholino)ethanesulfonic acid (MES) buffer pH 5.5. Subsequently, 0.2 mg of ethyl-3- (3-dimethylaminopropyl)-carbodiimide (EDC) and 0.5 mg of sulfo-N-hydroxysuccinimide (Sulfo-NHS) in 100 μL of MES were added and mixed for 15 min.

The excess of EDC/Sulfo-NHS was subsequently removed by centrifugation at 16,100g for 10 min. The resulting pellet was dissolved in phosphate buffered saline (PBS) pH 7.4. The final solution was stirred overnight and centrifuged again the next day, at 16,100× g for 10 min. Then, the pellet was resuspended in milli-Q water and stored at 4 °C.

### Characterization of gold nanorods and Chinese ink

The morphology and particle size of SPIONs was investigated by scanning transmission electron microscopy (STEM, FEI Quanta 250) operating at 10.00 kV. Dynamic light scattering system for nanoparticle analysis was used for size determination. The superficial charge was measured by a Zeta Potential Analyzer. Concentrations obtained from the synthesis were measured using Nanoparticle Tracking Analysis by NanoSight. Chinese ink was obtained from commercially available ARTEL and used at a concentration of 10%.

### Experimental design for retrograde and anterograde directional flow evaluation in mice

For *post-mortem* anterograde evaluations different nanoparticle solutions (gold nanorods and Chinese ink) were injected into the cisterna magna of C57BL/6 mice (n=3 per condition), after euthanasia by ketamine (300mg/kg) and xylazine (30 mg/kg) overdose. These were compared with control mice with no injections (n=3). A syringe with a 30G needle was loaded with 50 μL of each solution and administered in the cisterna magna; by placing the mouse in prone position, flexing the head at a 135^0^ angle with the body, and penetrating directly underneath and laterally to the end of the occipital bone towards the foramen magnum through the intact skin. The head and neck were preserved by fixation with 4% paraformaldehyde for histological Gold Enhancement (Nanoprobes GoldEnhance TM LM Kit) analysis.

For *post-mortem* retrograde flow evaluation different nanoparticle solutions (gold nanorods and Chinese ink) were injected into the deep cervical lymph node of C57BL/6 mice (n=3 per condition), after euthanasia by intraperitoneal ketamine (300mg/kg) and xylazine (30 mg/kg) overdose. These were compared with control mice with no injections (n=3). A syringe with a 30G needle was loaded with 50 μL of each solution and administered into the deep cervical lymph node. To locate the lymph node, skin and subcutaneous tissue were dissected at the midline of the neck, extending the field laterally at the supraclavicular area until both mandibular glands were exposed. Glands were detached from the clavicle surface and moved cranially. The sternocleidomastoid muscles were then displaced until the deep cervical nodes, surrounded by adipose tissue, were identified. The head and neck were preserved by fixation with 4% paraformaldehyde for histological Gold Enhancement (Nanoprobes GoldEnhance TM LM Kit) analysis.

For *in vivo* anterograde flow evaluation different nanoparticle solutions (SPIONs, exosomes loaded with SPIONs, gold nanorods, and Chinese ink) were injected into the cisterna magna of C57BL/6 mice (n=3 per condition). These were compared with control mice with no injections (n=3). Animals were anesthetized with 5% isoflurane and kept under anesthesia with a nasal cannula supplying 1% isoflurane during the entire procedure. A syringe with a 30G needle was loaded with 10 μL of each solution and administered in the cisterna magna; by placing the mouse in prone position, flexing the head at a 135^0^ angle with the body, and penetrating directly underneath and laterally to the end of the occipital bone towards the foramen magnum through the intact skin. Euthanasia by intraperitoneal sodium thiopental overdose (100 mg/kg) was performed 30 min after injection. The head and neck were preserved by fixation with 4% paraformaldehyde for MRI and histological analysis.

For *in vivo* retrograde flow evaluation different nanoparticle solutions (SPIONs, exosomes loaded with SPIONs, gold nanorods, and Chinese ink) were injected into the deep cervical lymph node of C57BL/6 mice (n=3 per condition; Chinese ink n=4). These were compared with control mice with no injections (n=3). Animals were anesthetized with 5% isoflurane and kept under anesthesia with a nasal cannula supplying 1% isoflurane during the entire procedure. A syringe with a 30G needle was loaded with 10 μL of each solution and administered into the deep cervical lymph node. To locate the lymph node, skin and subcutaneous tissue were dissected at the midline of the neck, extending the field laterally at the supraclavicular area until both mandibular glands were exposed. Glands were detached from the clavicle surface and moved cranially. The sternocleidomastoid muscles were then displaced until the deep cervical nodes, surrounded by adipose tissue, were identified. Euthanasia by intraperitoneal sodium thiopental overdose (100 mg/kg) was performed 30 min after injection. The head and neck were preserved by fixation with 4% paraformaldehyde for MRI and histological analysis.

The image acquisition was performed with a clinical Philips Achieva 1.5T MR scanner (Philips Healthcare, Best, Netherlands) and a single-loop surface coil (diameter=47 mm). Perls’ Prussian blue was used for iron tissular content analysis. Gold Enhancement (Nanoprobes GoldEnhance TM LM Kit) was used for GNR and Chinese ink analysis.

### Formalin-fixed, paraffin-embedded (FFPE) tissue processing for histology and special stains

Whole brain and neck samples were fixed for 24 hours on 4% PFA and then processed for paraffin embedding. Coronal sections of 4 μm were cut from each paraffin block, then sections were dried, deparaffinized and re-hydrated on distilled water. Gold Enhancement was performed with Nanoprobes GoldEnhance TM LM Kit according to manufacturer’s instructions. Once this procedure was done, the sections were counterstained with eosin. Nanoparticles were illustrated by Perls’ Prussian blue staining for iron content^25^. Tissue was deparaffinized and hydrated with distilled water, immersed in 10% aqueous potassium ferrocyanide and 20% aqueous hydrochloric acid for 20 min. Images of the stained slides were taken with an ICC50W Camera on a DM500 Leica Microscope at 4x, 10x, 20x, and 40x magnification.

Immunohistochemistry analysis was performed on the different sections with joint staining with Gold Enhancement and Perls’ Prussian blue to evaluate colocalization with an endothelial marker of lymphatic vessels (LYVE-1). After deparaffinization, antigen recovery was done using buffer Tris-EDTA pH 9.0 in a pressure cooker for 20 min. Endogenous peroxidase was blocked with 3% hydrogen peroxide for 15 min. Blocking of non-specific binding was performed with 3% BSA/PBS for 30 min. Overnight incubation at 4°C with the primary antibody, Recombinant Anti-LYVE1 antibody [EPR21771] (ab218535), was done at 1:5000 dilution. This was followed by incubation with the secondary antibody, Goat Anti-Rabbit IgG H&L (HRP) ab6721 (Abcam), for 1 hour at 25°C. Each slide was then developed with DAB for 1 min. Lung, gall bladder, and spleen tissues were used as controls. Quantification of histological images were performed with the software Image J.

### Animals

All procedures complied with regulations of the Research Ethics Committee of the Pontificia Universidad Católica de Chile. 49 Male C57BL/6 mice were purchased from the animal facility of the Pontificia Universidad Católica de Chile and housed in temperature and humidity-controlled rooms, maintained on a 12h/12h light/dark cycle.

Only adult animals (eight to ten weeks) were used in this study. Nine animals were assigned to *post-mortem* anterograde directional flow experiments, nine to *post-mortem* retrograde directional flow analyses, 15 to *in vivo* anterograde directional flow experiments, and 16 to the *in vivo* retrograde directional flow group. The sample size was chosen following similar, previously published research ^4,20,26,27^. Animals from different cages in the same experimental group were selected to assure randomization.

### Ethical approval

All experimental protocols were approved by the Research Ethics Committee of the Pontificia Universidad Católica de Chile, the CEC-CAA (Comité Ético Científico para el Cuidado de Animales y Ambiente), with Protocol ID: 190826005. This study was conducted according to ARRIVE guidelines.

## Acknowledgements

We thank Sergio Alvarez for his collaboration with electroporation equipment. The authors acknowledge Wilda Olivares, Andrés Rodriguez, Pablo Santoro, Ignacio Wichmann, Rocío Bustos, Leticia Gonzalez, Miguel Urrutia, Mauricio Cuello, and Flavia Zacconi for comments and assistance. This study was supported by ANID BECAS/DOCTORADO NACIONAL 21211334; ANID - Millennium Science Initiative Program - ICN2021_004, ANID – FONDECYTs 1220922, 1231773 and 1211482; CONICYT-FONDAP 15130011, FONDEQUIP NTA (EQM160157), and FONDEQUIP SEM (EQM170111).

## Author contributions

H.M.R.Z. and I.P. conceived the study. H.M.R.Z., I.P., E.S.H. A.H.C., M.J.K., C.P.Y., J.E.O., A.N.T., P.V.B., and M.E.A. designed the experiments. H.M.R.Z., I.P., E.S.H., P.J.G., A.N.T., P.V.B., V.A.C., E.A.M., and A.R. performed preparation and characterization of nanoparticles. H.M.R.Z., C.P.Y., and C.M. conducted mouse experiments. D.S. performed histological analysis and immunohistochemistry techniques. J.E.O. conducted cytotoxicity analysis. All authors participated in data analysis. H.M.R.Z. wrote the paper and all the authors contributed to its editing.

## Data availability

The data that support the findings of this study are available from the corresponding author upon reasonable request.

## Corresponding author

Correspondence and requests for materials should be addressed to Héctor M. Ramos-Zaldívar. Primary.

---

**Supplementary Fig. S1:**
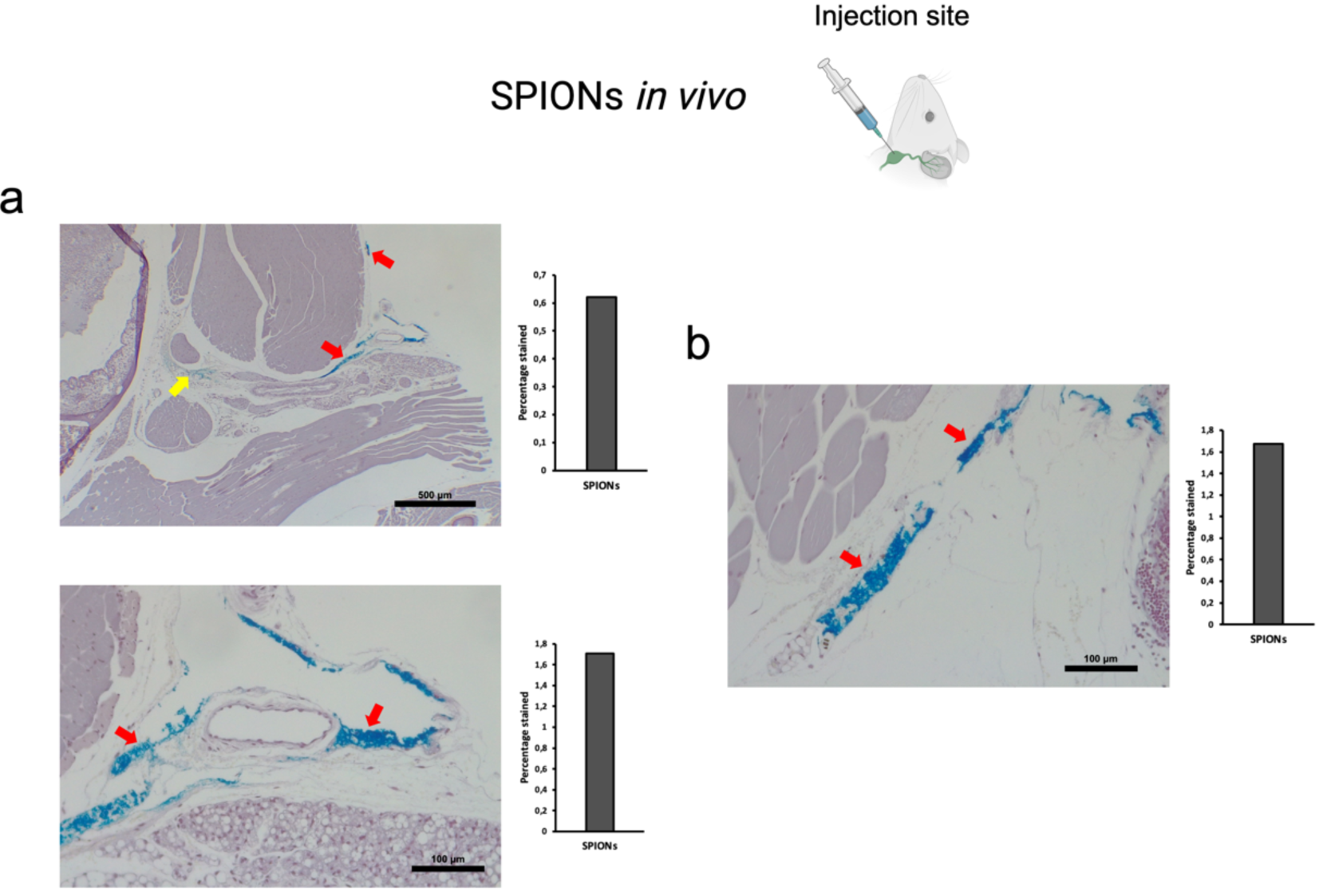
Retrograde directional flow analysis after *in vivo* administration of SPIONs into the deep cervical lymph node. **a**, Perls’ Prussian Blue shows staining of lymphatic vessels of the head (red arrows), including the meningeal lymphatic vessels, as well as connective tissue (yellow arrow) (n=3). **b**, SPIONs were also identified within lymphatic vessels of the neck (red arrows) (n=3). SPIONs: superparamagnetic iron oxide nanoparticles.

**Supplementary Fig. S2:**
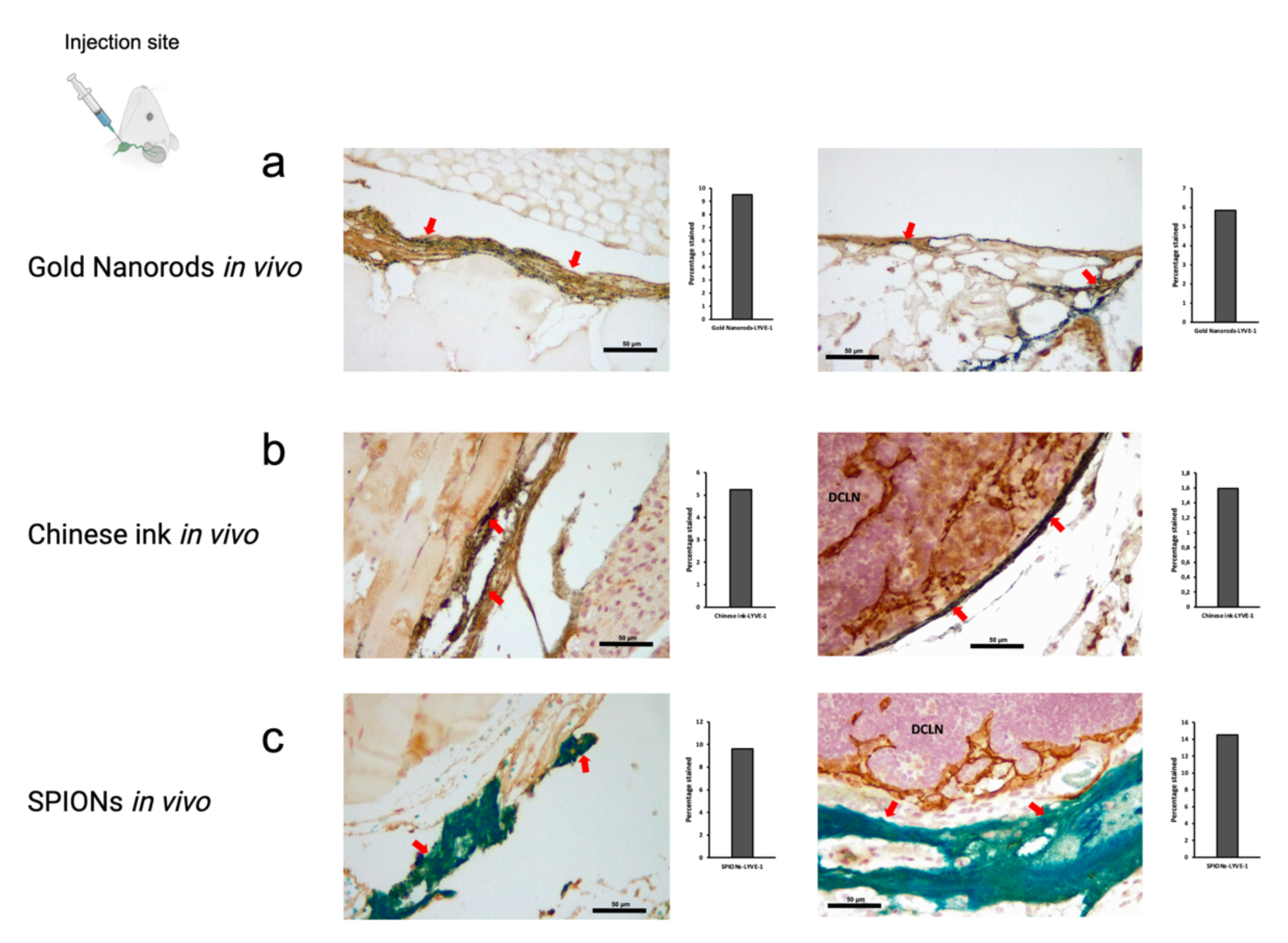
Retrograde directional flow analysis of cervical structures after *in vivo* administration of nanoparticles into the deep cervical lymph node. **a**, Combined Gold Enhancement and anti-LYVE-1 immunohistochemistry showed gold nanorods within lymphatic vessels of the neck (red arrows) (n=3). **b**, Anti-LYVE-1 immunohistochemistry revealed Chinese ink nanoparticles within cervical lymphatic vessels and lymph nodes (n=4). **c**, Combined Perls’ Prussian Blue and anti-LYVE-1 immunohistochemistry showed SPIONs within cervical lymphatic vessels and lymph nodes (n=3). LYVE-1: lymphatic vessel endothelial hyaluronan receptor-1; SPIONs: superparamagnetic iron oxide nanoparticles.

**Supplementary Fig. S3:**
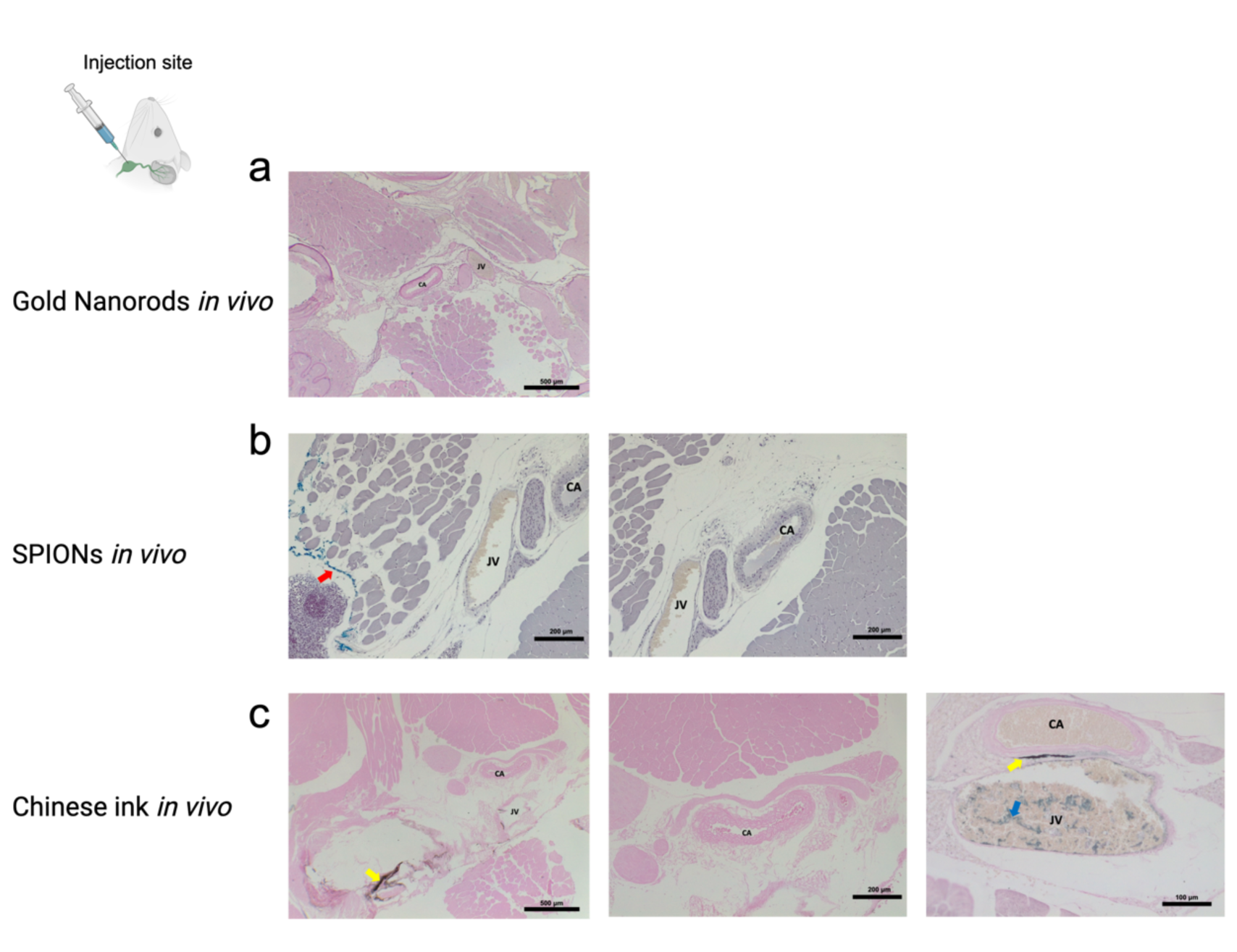
Histological analysis of arterial and venous vessels after *in vivo* administration of nanoparticles into the deep cervical lymph node. **a**, No Gold Enhancement staining was identified within the carotid arteries or jugular veins after gold nanorod injections in all mice (n=3). **b**, No Perls’ Prussian Blue staining was detected within the carotid arteries or jugular veins after SPIONs administration (n=3). **c**, One mouse presented staining for Chinese ink within the jugular vein but not the carotid arteries. The other three animals did not show staining of arterial or venous cervical structures. SPIONs: superparamagnetic iron oxide nanoparticles.

**Supplementary Fig. S4:**
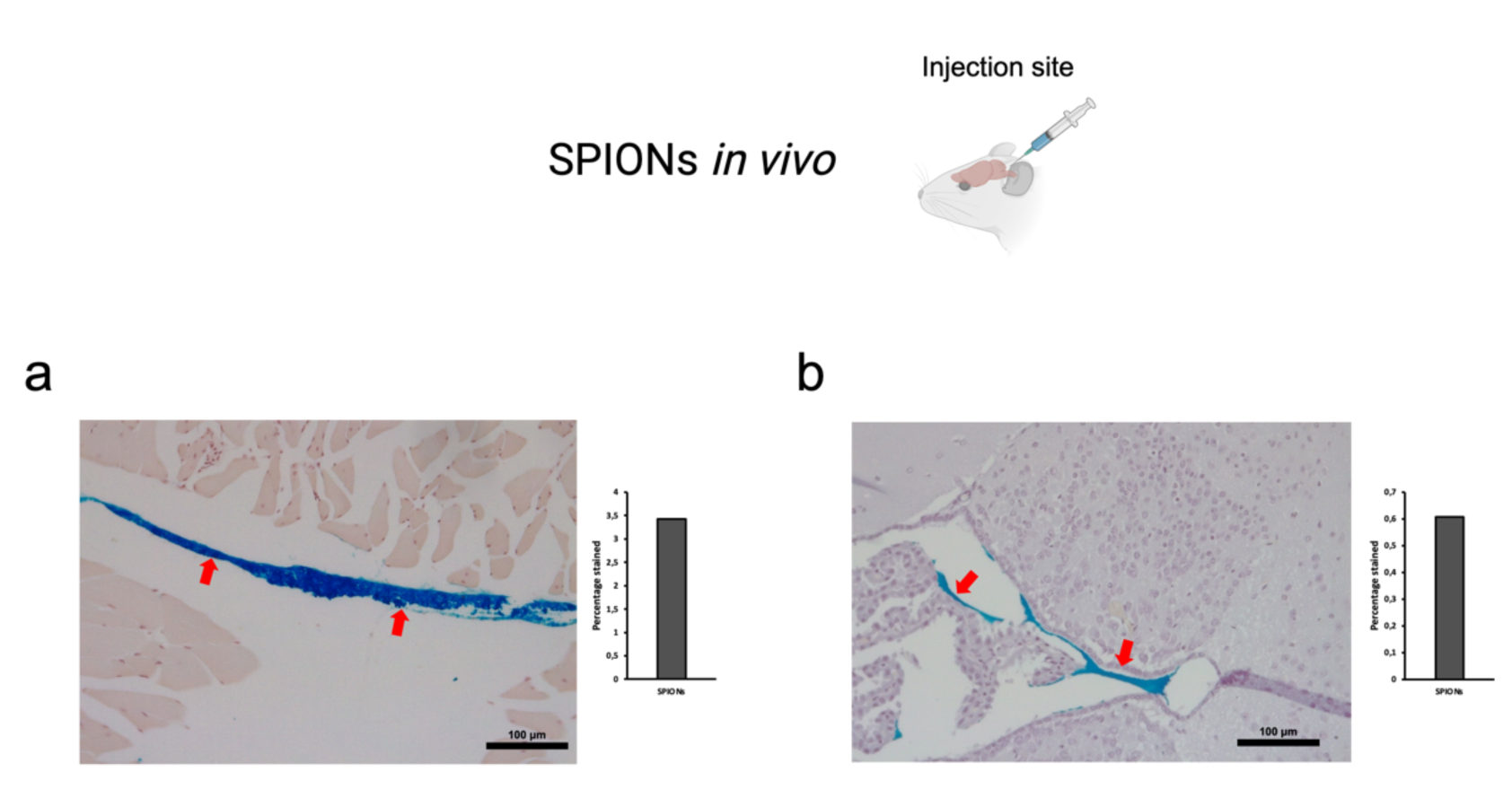
Anterograde directional flow analysis after *in vivo* administration of SPIONs into the cisterna magna. **a**, Perls’ Prussian Blue shows staining of cervical lymphatic vessels (red arrows) (n=3). **b**, SPIONs were identified within ventricular spaces of the brain (n=3). SPIONs: superparamagnetic iron oxide nanoparticles.

